# Quantitative analysis of 3D alignment quality: its impact on soft-validation, particle pruning and homogeneity analysis

**DOI:** 10.1101/088062

**Authors:** J. Vargas, R. Melero, J. Gómez-Blanco, J. M. Carazo, C. O. S. Sorzano

## Abstract

Single Particle Analysis using cryo-electron microscopy is a structural biology technique to capture the three-dimensional conformation of biological macromolecules. The projection images used to construct the 3D density map are characterized by a very low signal-to-noise ratio to minimize radiation damage in the samples. As a consequence, the 3D alignment process is a challenging and error prone task and this job usually determines the success or failure of the macromolecule reconstruction. In this work, we present a soft-alignment validation approach, which can quantify the alignment precision and accuracy as well as the data homogeneity of the single particles when they are confronted with the resultant reconstructed 3DEM map. We have also applied this method to data homogeneity analysis and particle pruning, improving the data quality and as a consequence the final map resolution.

## Introduction

Understanding how macromolecular complexes fulfill their complicated roles in the living cell is a central theme in molecular biology. Structural biology aims to deduce how such complexes function by determining the 3D arrangement of their atoms. Several techniques may be used to determine such structures. By far, the most successful technique has been X-ray crystallography. Assuming that the macromolecular complex of interest can be crystallized, this technique may yield atomic resolution and is not limited by the size of the complex. Nuclear magnetic resonance may provide unique information about dynamics and interactions, but atomic structure determination is restricted to small complexes; that is, those with molecular weights below 40-50 kDa. Both techniques typically require large amounts of relatively pure sample (on the order of several mg).

Single Particle Analysis (SPA) is a form of cryo-electron microscopy, which allows obtaining three-dimensional information of both large and small macromolecules close to their native state, captured in the process of performing their work, and using sample preparations with low concentrations of samples (0.1 mg/ml of purified sample may be enough) [Wang2006; Frank2009]. The idea of data collection in single particle analysis is to assume that the macromolecule or particle occurs in multiple copies with (essentially) identical structure, and that its orientation samples the entire angular range without leaving major gaps. Thus, instead of having to tilt the grid on which the sample is spread into multiple angles (as in electron tomography), it is then possible to take snapshots of multiple molecular views and, after suitable alignment and image processing, combine all projections into a density map depicting the molecule in three dimensions. To perform this image processing step, there are different software packages as Xmipp [DelaRosa-Trevín2013], Eman [Ludtke1999] or Relion [Scheres2012], among others.

At present, one of the main problems of this technique is that there are not extensively used and accepted validation methods in SPA to assess the validity and quality of a reconstructed density map. A recent controversy highlighted this issue [Mao2013a; Mao2013b; Henderson2013; Subramaniam2013; vanHeel2013], imposing high priority to the development of novel methodologies to address this limitation [Henderson2012]. Currently, the only way to perform a quantitative validation of a 3DEM map requires the analysis of pairs of single particle images recorded at different tilt angles (tilt-pairs) [Henderson2011]. The tilt-pairs validation method works comparing the discrepancy between the calculated orientations among non-tilted and tilted particles, with respect to the known tilting angle. This discrepancy is a good indicator of the 3D map quality; nevertheless collecting high-resolution and high-quality tilt-pairs is itself a relatively challenging process as often drift and/or charging occur in the tilted images [Murray2013]. Furthermore, for small cryogenized samples the determination of particle correspondences between the untilted and tilted micrographs can be a difficult and error-prone process. In many occasions, the map evaluation is performed retrospectively and then tilt pairs may not be available. Additionally, this approach requires increasing the amount of data to collect and process. Finally, beam-induced movement may introduce an extra uncontrolled source of inconsistency or dispersion in the tilt-pair plot. These issues explain that although this test is currently the only fully accepted map validation methodology, it has not been extensively adopted by electron microscopy practitioners. Recently, new methods to determine map quality scores have been proposed [Heymann2014; Stagg2014; Vargas2016]. In [Heymann2014] an approach is proposed to detect “the phantom in the noise problem” meaning reconstructed density maps computed from pure noise images aligned to that map. In this case, alignment of a limited number of particle and noise images against the final released map —used as reference— is required. Finally, the resolution for the noise and particle maps is obtained through FSC analysis by goldstandard procedure, and further analysis of the resolutions achieved from particle and pure noise maps is done. In [Stagg2014] it is proposed an empirical quality metric for cryo-EM reconstructions by a ResLog plot. This representation shows the inverse of the resolution achieved versus the logarithm of the number of particles used, and provides heuristic information about the consistency between the obtained map and the particle dataset. Finally, in [Vargas2016] it is proposed an approach to determine the alignment precision of a set of particles used in the map reconstruction process. This information is useful to determine the particle alignment reliability with respect to the reconstructed structure and can be used to define a map quality score. The main limitations of this approach are: 1) this method can determine the alignment precision for each particle but not its alignment accuracy, hence it cannot be considered a proper alignment validation approach. Note that precision refers to the reproducibility and repeatability of a measure, while accuracy is related to the degree of closeness of an obtained quantity with respect to the true value; 2) there are map projections which have more capacity to be aligned with precision than others. Note that the alignment precision highly depends on how the energy is distributed in the images. As a consequence, particles may not be ranked according to this score; 3) the particle alignment precision is determined using only one reference, which corresponds to an uniform distribution of random angular directions within the asymmetric unit, which may not be a sufficiently demanding requirement.

In this work, we propose an approach that provides both the alignment precision and alignment accuracy for each particle used in the reconstruction. These scores can be used to soft-alignment validation of a 3DEM map with respect to the set of particles used in the reconstruction process. The percentage of particles that aligns with both precision and accuracy is an indicator of the goodness of a reconstruction. Trustworthy reconstructions should provide high percentage values for these scores. In addition, these indicators provide information about the homogeneity of the data. Nonhomogeneous data will not be able to provide high-resolution reconstructions. In order to improve the data homogeneity, we propose to use these alignment precision and accuracy scores to prune the particles. The term soft-alignment validation refers to necessary conditions that a valid 3DEM map may verify but it does not stand for sufficient conditions for a map to be correct. This means that if a 3DEM structure provides a low soft-alignment validation score this map should not be accepted. On the other hand, maps with high scores can still represent incorrect structures, but still consistent with the data (false positives).

## Results

### Outline of the method

The goal of this method is to provide objective information about the alignment precision and accuracy for each experimental particle used in the reconstruction. The input of the method is a set of experimental particles previously aligned by any method and the corresponding reconstructed map. The approach is based on three steps that should be run for each particle: alignment precision estimation, alignment accuracy estimation, and determination of the percentage of reliable particles (*Q* value).

#### Particle precision estimation

Determination of the alignment precision of each particle is achieved analyzing the orientation distribution of the most similar map projections in the unit projection sphere and within the asymmetric unit. The map gallery of images is obtained projecting the volume into a regular grid of orientations, with a typical angular sampling rate of 5°. The more clustered is this orientation distribution, the higher the angular precision for the particle is. The clusterability of this distribution of orientations may be quantified using the Hopkins clustering tendency parameter [Banerjee2004], as was previously done in [Vargas2016]. However, this descriptor is insensitive to cases where the orientation distribution forms two or more different clusters. In this case, we have quantified the orientation clusterability of particle *m* by the following functional:

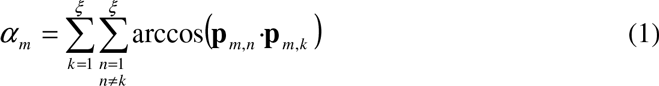

with *ξ* the number of most similar map projections (usually *ξ*=6), ‘·’ refers to the inner product and **p***_m,n_* the projection direction coordinates of the *n*th most likely projection directions of particle *m*. Observe that *α_m_* is sensitive to cases with two or more different clusters. This cluster tendency parameter provides an alignment precision value for each particle. To determine if this numeric value can be considered “good” or “bad”, references are required to compare it to.

A “bad” reference can be defined from a random uniform orientation distribution within the asymmetric unit. We compute the corresponding clusterability parameter *α_NOISE_* using Expression (1) from ***ξ*** random angular assignments defined in the projection sphere and within the asymmetric unit. As ***ξ*** is usually small, we repeat this experiment *M* times (*M*∼500) obtaining finally a mean clusterability parameter ***α_NOISE_***.

To define a “good” reference we can construct a synthetic set of “perfect” particles totally compatible with the input volume. This set is constructed projecting the map at the same orientations and distorted by the same CTFs than the experimental projections. Therefore, for each experimental particle (denoted by *m* index), we compute its respective “perfect” counterpart. The clusterability parameter of this “perfect” projection set is determined following the same procedure explained before obtaining ***α****_m,good_*. In order to decide if the clusterability parameter ***α****_m_* of a experimental particle *m* refers to precise alignment, we should compare this value with the two obtained references 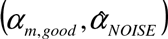. To this end, we map the clusterability scores 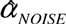 and *α_m,good_* to subjective quality precision alignment values of 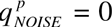 and 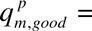 1 respectively. From these two points 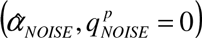 and 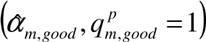 we can fit a straight line that pass through them and determine the quality precision alignment of experimental particle *m* as

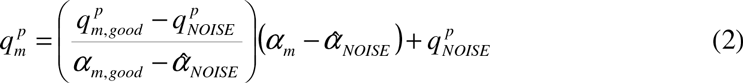

Observe that the defined quality precision alignment score is an easy to interpret parameter: the closer 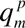 is to 1 (or to 0) the better (the worse) the alignment precision is. Therefore, we can establish as criteria that an experimental particle aligns with precision if 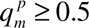, or any other threshold chosen by the user. Note that this threshold is arbitrary and it does not hold a direct statistical meaning. In Figure 1 we show a scheme of the alignment precision estimation process for each experimental particle.

**Figure 1.**
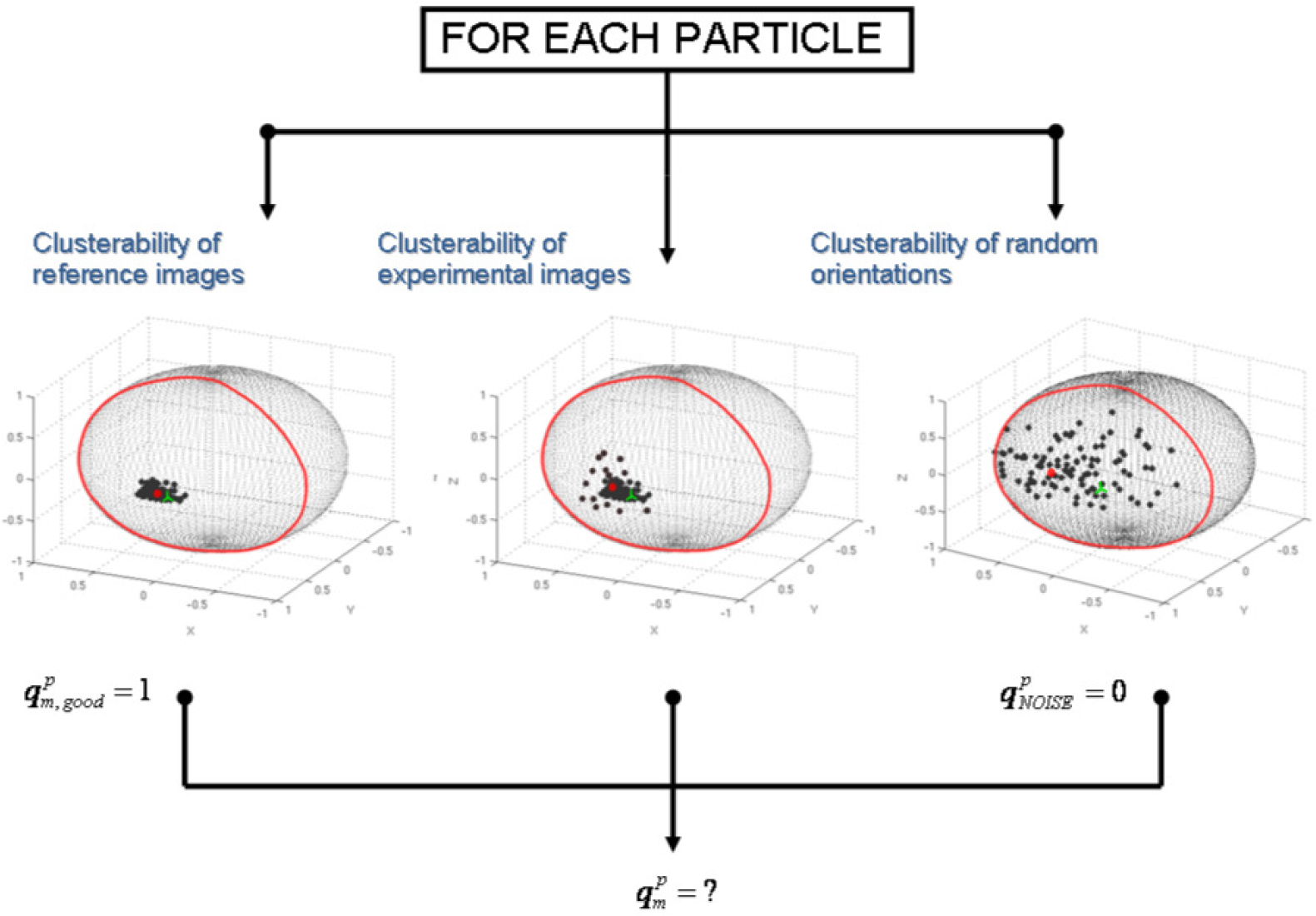
Diagram of the alignment precision and accuracy estimation process for each experimental particle. First, the clusterability of the orientation distribution is obtained for the experimental, perfect and pure noise particles. These values are used to give an alignment precision score to each experimental particle between 0 and 1. In addition, the alignment accuracy is computed comparing the previously obtained particle orientation (green cross) with the weighted average orientation of most similar map projections (red point).

#### Particle accuracy estimation

Determination of the alignment accuracy for each particle is performed comparing the previously computed particle orientation, used in the map reconstruction to be analyzed, with the weighted average orientation of the most similar map projections. The weights come from the cross-correlation between the particles and their most similar map projections, which were determined after a global alignment search using the input 3DEM map as reference. This comparison is quantified by the geodesic distance between these two orientations (previous and weighted average) in the projection sphere and it is denoted here by *χ_m_*. Conceptually, here we are comparing the final computed orientation by the reconstruction method after refinement with the orientation obtained from a pure global alignment search through cross-correlation and using as template the final three-dimensional map. Usual iterative reconstruction methods perform global particle angular searches only in the first iterations of the reconstruction process. These angular explorations transform to local searches as the number of iterations increase. This means that the alignment process of reconstruction approaches is generally very dependant on previous decisions taken, with the risk of getting trapped into an alignment local minima for a given particle. Hence, if the alignment approach makes an erroneous angular assignment for one particle at any iteration, this incorrection will not be amended at following iterations.

The accuracy parameter obtained for each particle (*χ_m_*) is difficult to interpret, as this geodesic distance depends on parameters as the symmetry and angular sampling, among others. In order to decide if a particle aligns with accuracy, we need two references to compare it to. As before, we can use the synthetic “perfect” particle set projecting the map at the same orientation and distorted by the same CTF than the experimental projections but without added noise. Therefore, for each experimental projection *m* we can determine a “good” reference accuracy parameter (*χ_m, good_*). On the other hand, a “bad” reference can be determined comparing the previously computed orientation by the reconstruction method with the orientation obtained by a weighted average of *N* random orientations within the asymmetric unit. As before, this comparison is performed *M* times obtaining finally a mean accuracy parameter 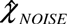. From these two references, we determine the quality accuracy alignment score for each experimental particle *m* as

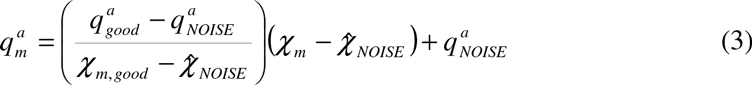

where 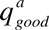 and 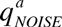 equals to 1 and 0, respectively. We can establish as criteria that an experimental particle aligns with accuracy if 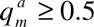 (or by any other threshold selected by the user). In Figure 2 we show a scheme of the typical alignment procedure, which starting from a low-resolution initial model performs a global angular search for each particle (a). This global search converts to a local one as the number of iterations and the map resolution increases (b-d). Figure 2(b) exemplifies the case when an incorrect angular assignment can be made for one particle. In Figure 2(e) we show how the particle alignment accuracy is determined by a global angular search using the final reconstructed map.

**Figure 2.**
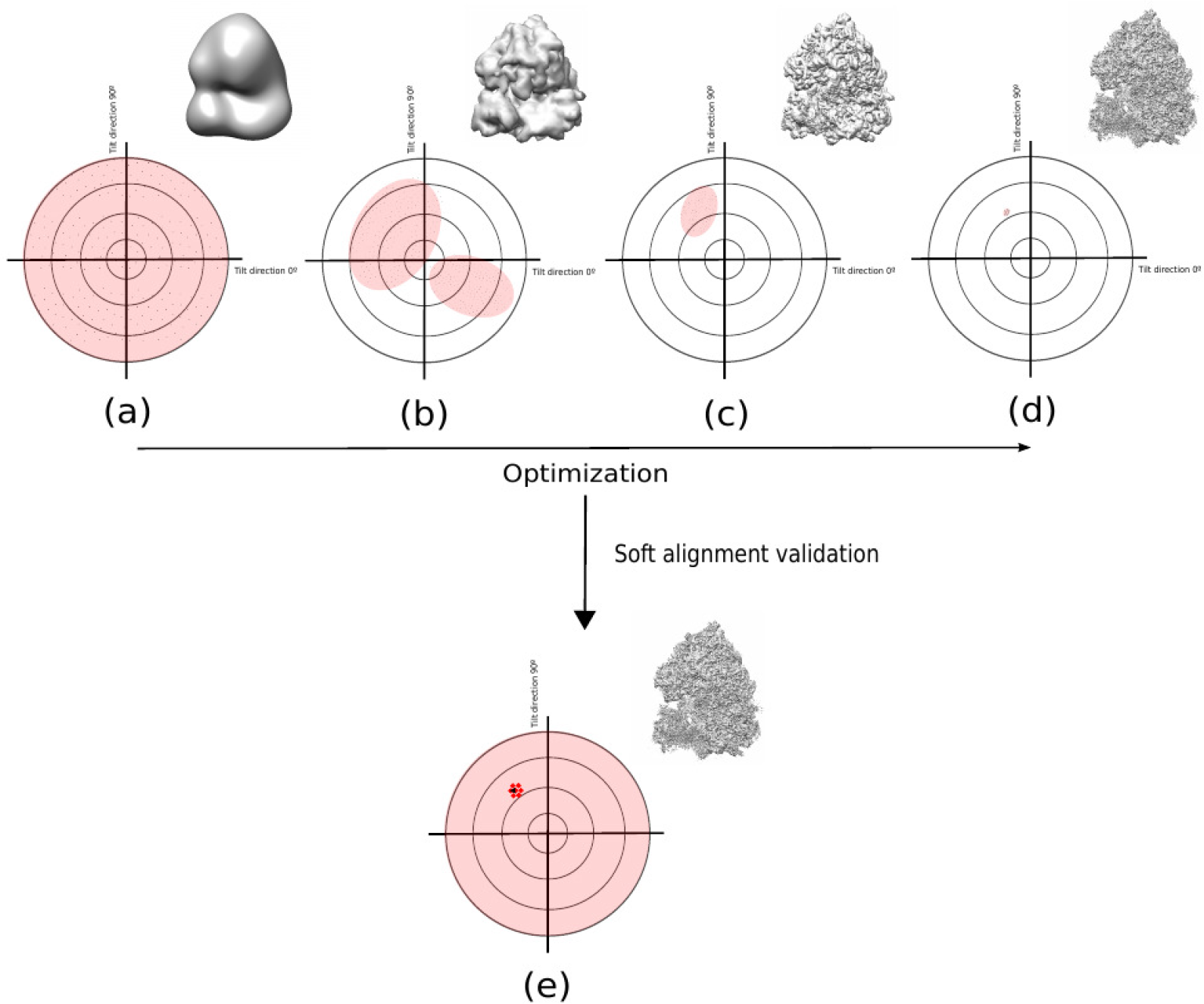
Typical alignment procedure performed for each particle. Starting from a low-resolution initial model the alignment method performs a global angular search with low angular sampling (a). This global search converts to a local one (increasing the angular sampling) as the number of iterations and the map resolution increases (b-d). Our proposed approach angular accuracy estimation is based on re-computing the particle orientation by a global search using the final reconstructed map (e).

#### Determination of the percentage of reliable particles

After we have obtained for each experimental particle the quality precision and accuracy parameters 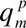 and 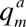, we can determine a global alignment parameter denoted as *Q*, which determines the percentage of particles that aligns with both precision and accuracy

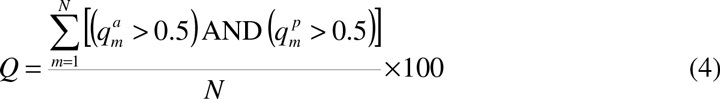

with *N* the number of particles in the dataset. Obviously *Q* gives information about the consistency between the 3DEM map and the experimental particles. The larger *Q* is, the better the consistency between the particles and the 3DEM map.

### Application examples

We have used the proposed approach in different cases, including soft-validation analysis of high-quality (β-galactosidase) and low quality (HIV-1 Env trimer) maps and application to a heterogeneous dataset (Ribosome).

#### First Experiment: Application to high-resolution data

We have applied our proposed approach to high-quality data corresponding to the β- galactosidase complex. We have used the data of the 2015 Map Challenge, with EMPIAR codes EMPIAR-10012 and EMPIAR-10013. These data was obtained with a FEI Titan KRIOS microscope and using a Gatan K2 detector. We used the initial particle coordinates given for the Map Challenge. A gold-standard approach was performed dividing the data in two independent halves, each composed by 11,412 particles. The final reconstructed 3DEM map, obtained by Relion [Scheres2012] through Scipion framework [DelaRosa2016], has a 0.143-FSC resolution of 3.25 Angstroms, and secondary structure elements can be seen clearly. This map and the resultant FSC are shown in Figure 3(a) and (b), respectively. In addition, we have also computed the FSC curve between the reconstructed map and the corresponding deposited PDB structure (PDB code: 3j7h). The resultant 0.143-FSC resolution is of 3.02 Angstroms. Using this data, we run our proposed soft-alignment validation approach to each half obtaining the results shown in Figure 4. In this figure each red point represents the alignment precision (x-axis) and accuracy (y-axis) for one experimental particle. Observe that these soft-alignment validation maps show a clear cluster around point (1, 1). The percentage of particles that align with precision, accuracy and both precision and accuracy are given in Table 1. As can be seen from Table 1, most of the particles align with precision and accuracy, being this an indicator of the map high quality. Additionally, observe that there is a significant percentage of particles which aligns with precision but not with accuracy. This indicates the presence of compositional, conformational or artefact-based heterogeneity, as bright spots for example in the dataset. The β-galactosidase complex is well known to not present conformational heterogeneity. We have visually checked that a significant amount of particles are of low quality and affected by artefacts. Visually, it is confirmed that most of the particles with low alignment accuracy or both accuracy and precision are affected by artefacts. As example, in Figure 5, we show a set of downsampled particles which show high precision but low accuracy in the first row and both high precision and accuracy in the second row. Taking into consideration that with the proposed method we can rank the particles according to its alignment quality, we have performed a particle pruning process. We have rejected particles showing values of alignment precision or accuracy lower than 0.5. As result, after pruning we have reduced the amount of particles approximately 21% and we have rejected 4,875 images from the set of 22,824 particles. We have reconstructed the density map from the selected set of 17,949 particles and the resulting 0.143-FSC resolution is of 3.00 Angstroms when this map is confronted with the corresponding PDB. In Figure 6, we show the resultant FSC curves obtained when the PDB is confronted with the pruned (red solid curve) and not pruned 3DEM maps (blue dashed curve). As can be seen from these curves the pruned map presents better quality than the other over all range of frequencies and using approximately 21% less data.

**Figure 3.**
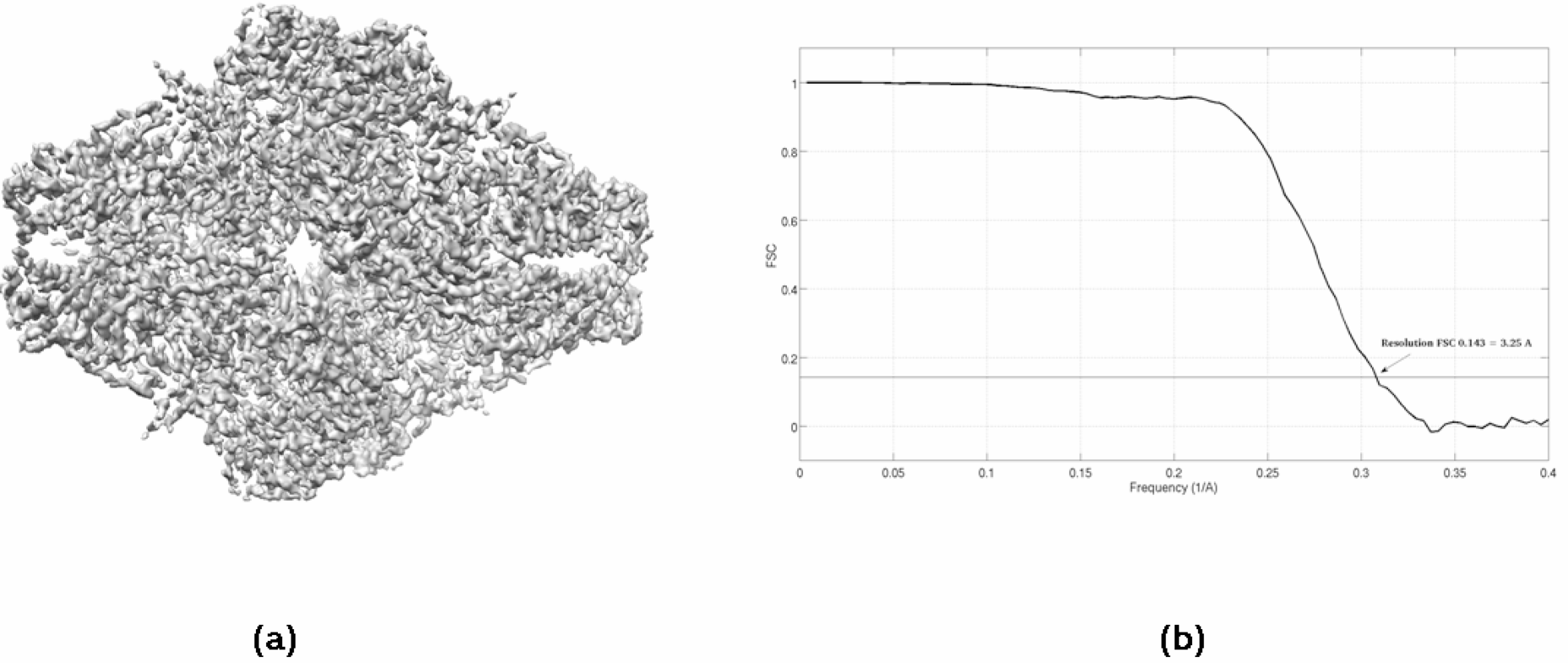
Reconstructed 3DEM map for the β-galactosidase complex of the 2015 Map Challenge with EMPIAR codes EMPIAR-10012 and EMPIAR-10013 (a) and obtained FSC curve based on Gold standard approach. The resultant 0.143-FSC resolution is of 3.25 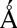.

**Figure 4.**
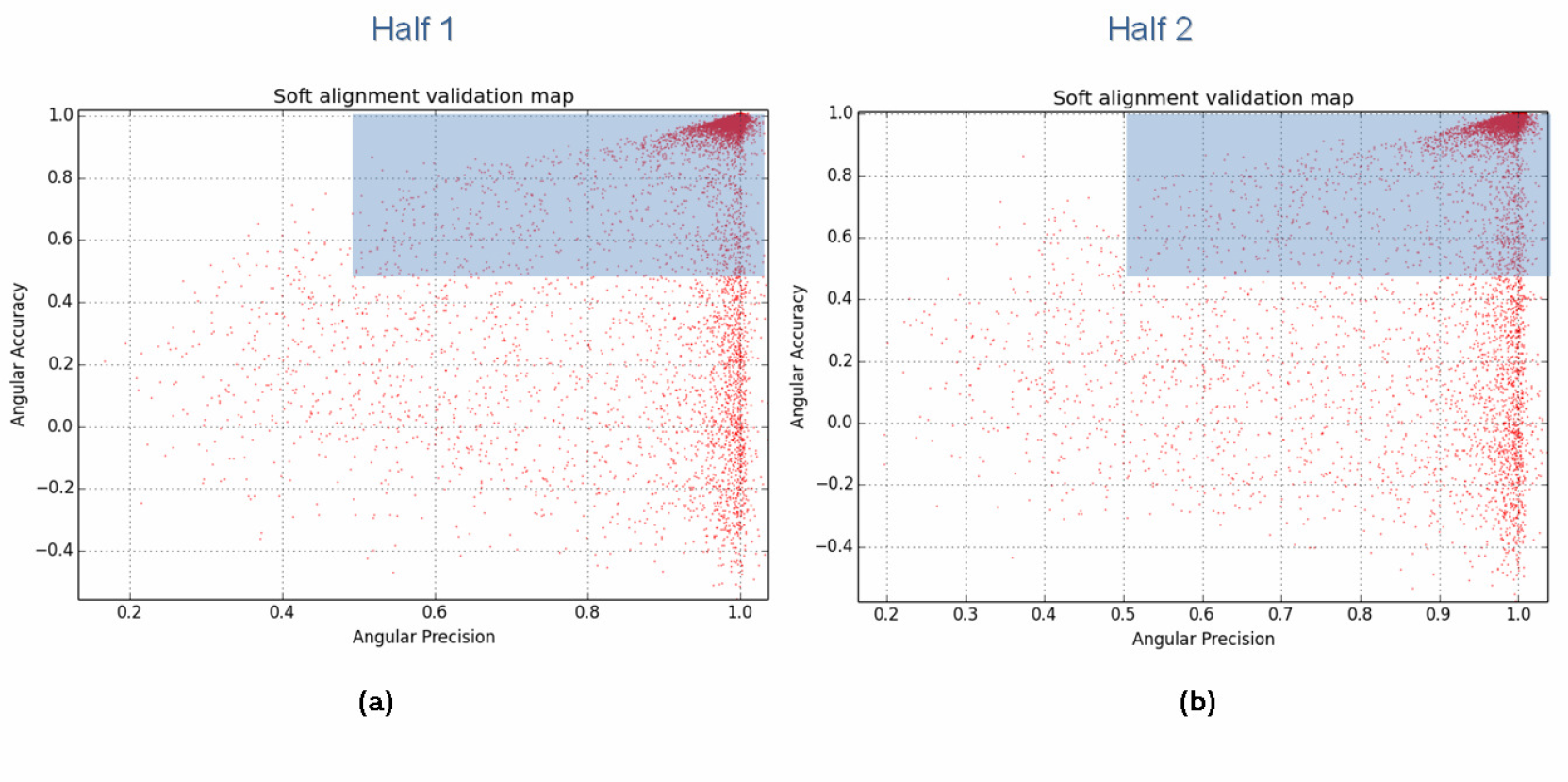
Soft-alignment validation maps for each half of the β-galactosidase complex and composed by 11,412 particles. The blue partially transparent rectangle indicates the particles which align with both accuracy and precision.

**Figure 5.**
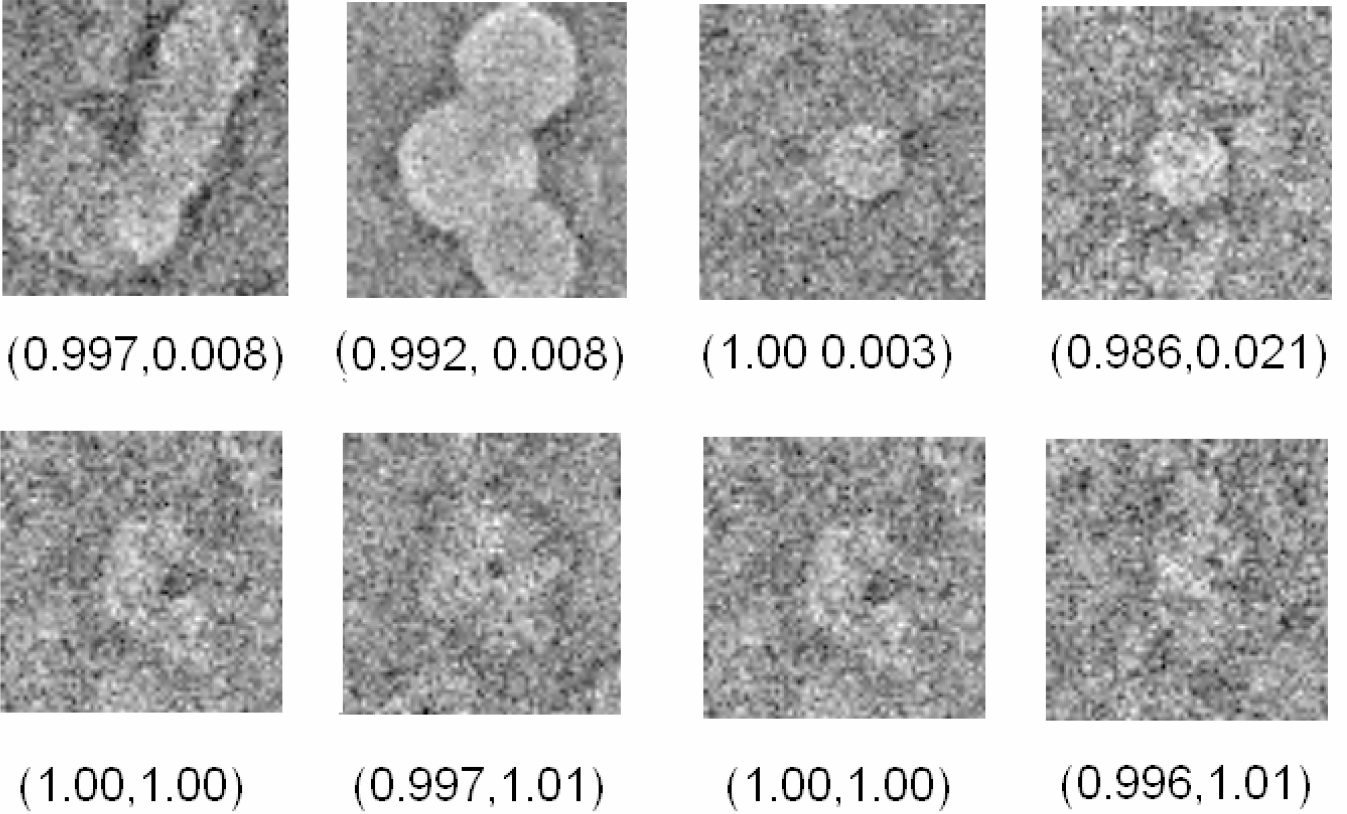
Exemplifying set of downsampled β-galactosidase particles showing high precision but low accuracy in the first row and both high precision and accuracy in the second row.

**Figure 6.**
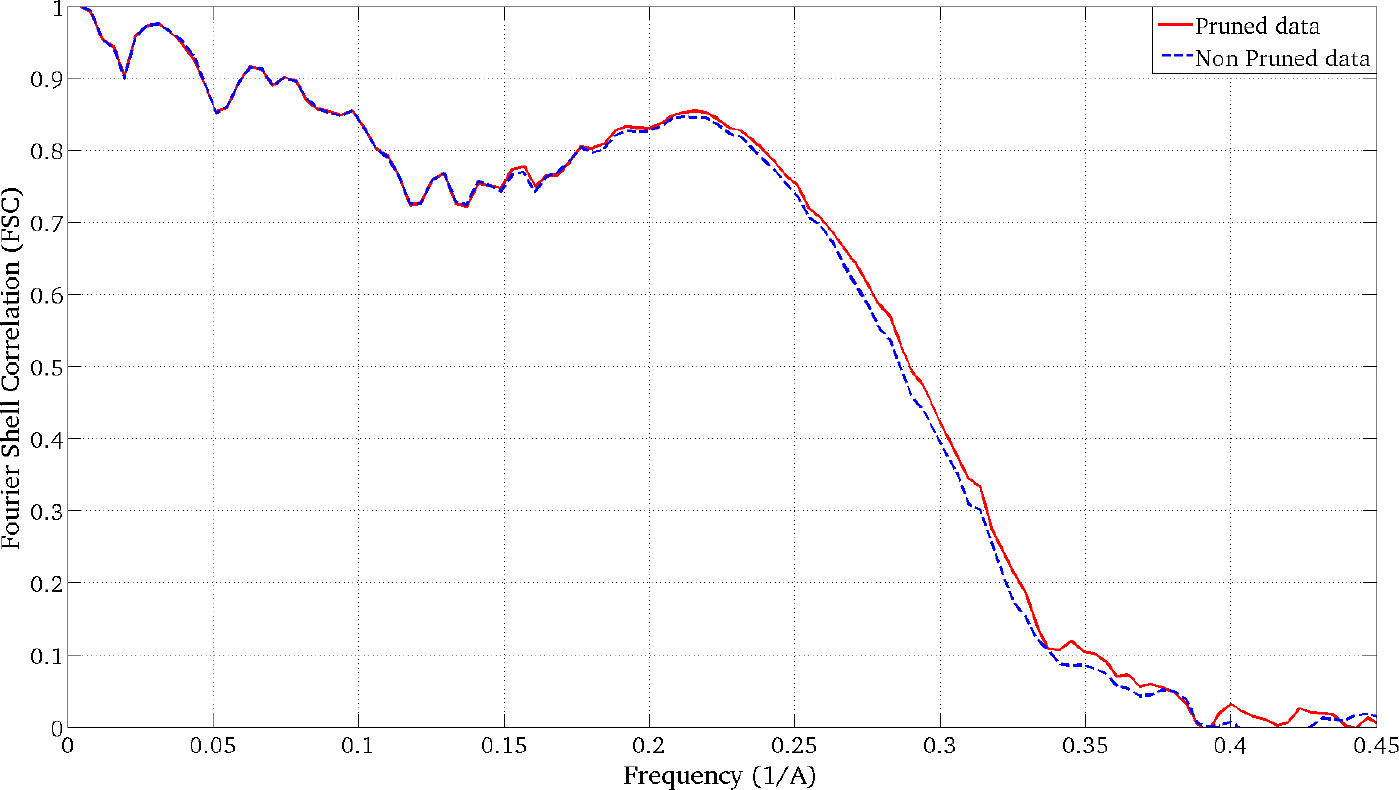
FSC curves obtained when the pruned (red solid curve) and non-pruned (blue dashed curve) 3DEM maps obtained for the β-galactosidase complex are confronted with the PDB map.

**Table 1.** Percentage of particles that align with precision, accuracy and both precision and accuracy for each half of the β-galactosidase complex.

|  | **Half 1** | **Half 2** |
| --- | --- | --- |
| **Precision (%)** | 97 | 97 |
| **Accuracy (%)** | 79 | 79 |
| **Precision & Accuracy (%)** | 79 | 79 |

#### Second Experiment: Application to ranking of *ab initio* initial maps

We have applied the soft-alignment validation approach to the task of ranking different *ab initio* initial maps. We have used as before the β-galactosidase complex. 2D classes were obtained from picked particles using CL2D method [Sorzano2010]. From these averages we run RANSAC *ab initio* initial volume estimation method [Vargas2015], obtaining ten different low resolution maps. From this set, we selected two maps, one clearly correct and another of lower quality. In Figure 7 we show the used 2D classes and the two initial volumes selected. The obtained results for the alignment precision when these two maps are confronted with 11,412 particles, composing each of the two halves of the full dataset, are 0.98 and 0.90 for maps shown in Figure 7(a) and (b), respectively. These values indicate that in both cases we can align the particles with precision but (a) map gives better results. Note that in this experiment we cannot determine the alignment accuracy as at the moment of obtaining the initial map, we have not yet aligned the particles. In Figure 8 we show the respective alignment precision plots obtained when the maps shown in Figure 7(a) and (b) are confronted with the particles. As can be seen from this figure the “correct” map presents higher alignment precision than the other one.

**Figure 7.**
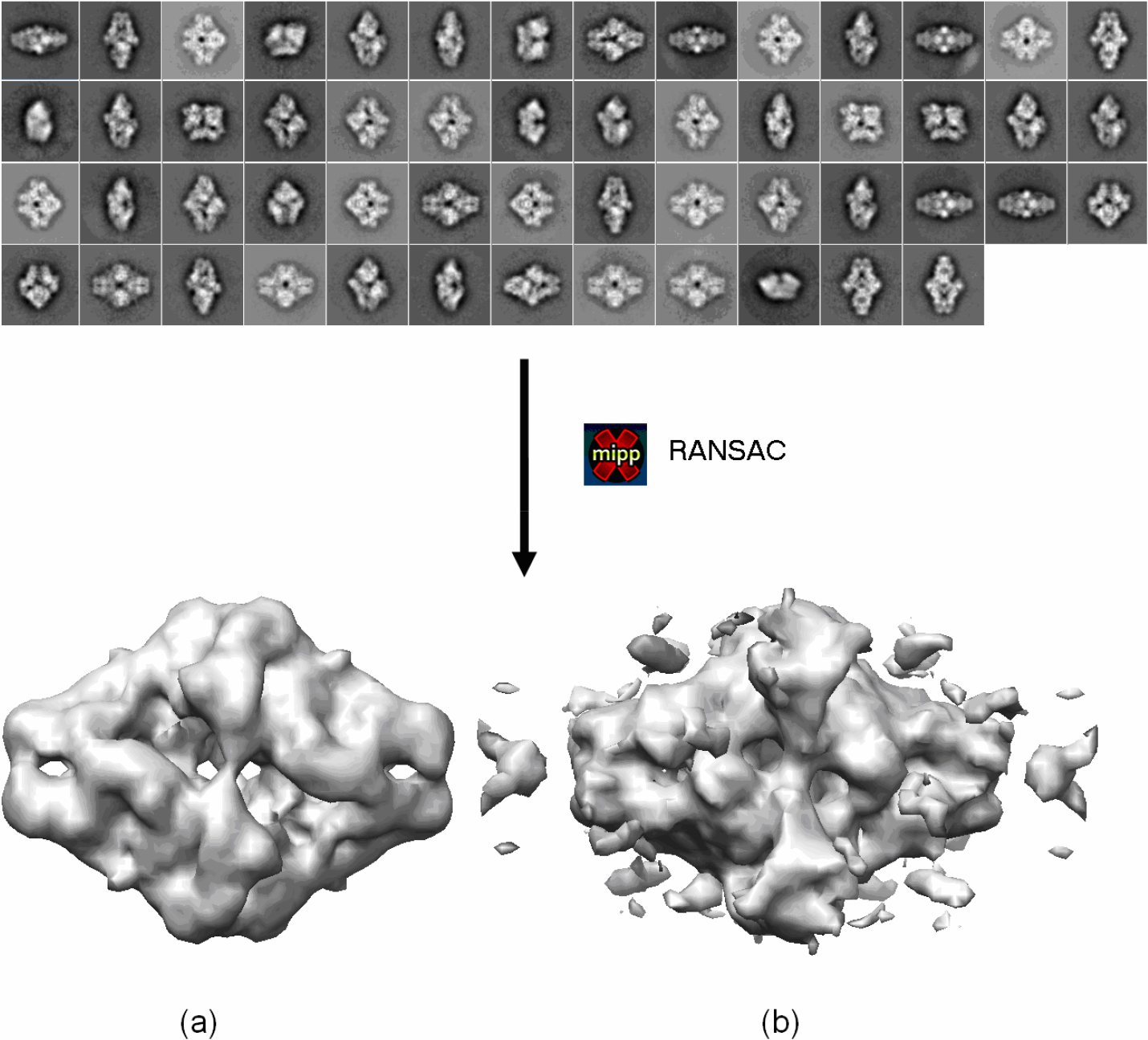
Experimental class averages of β-galactosidase particles and obtained “good” (a) and “bad” quality (b) *ab initio* initial volumes.

**Figure 8.**
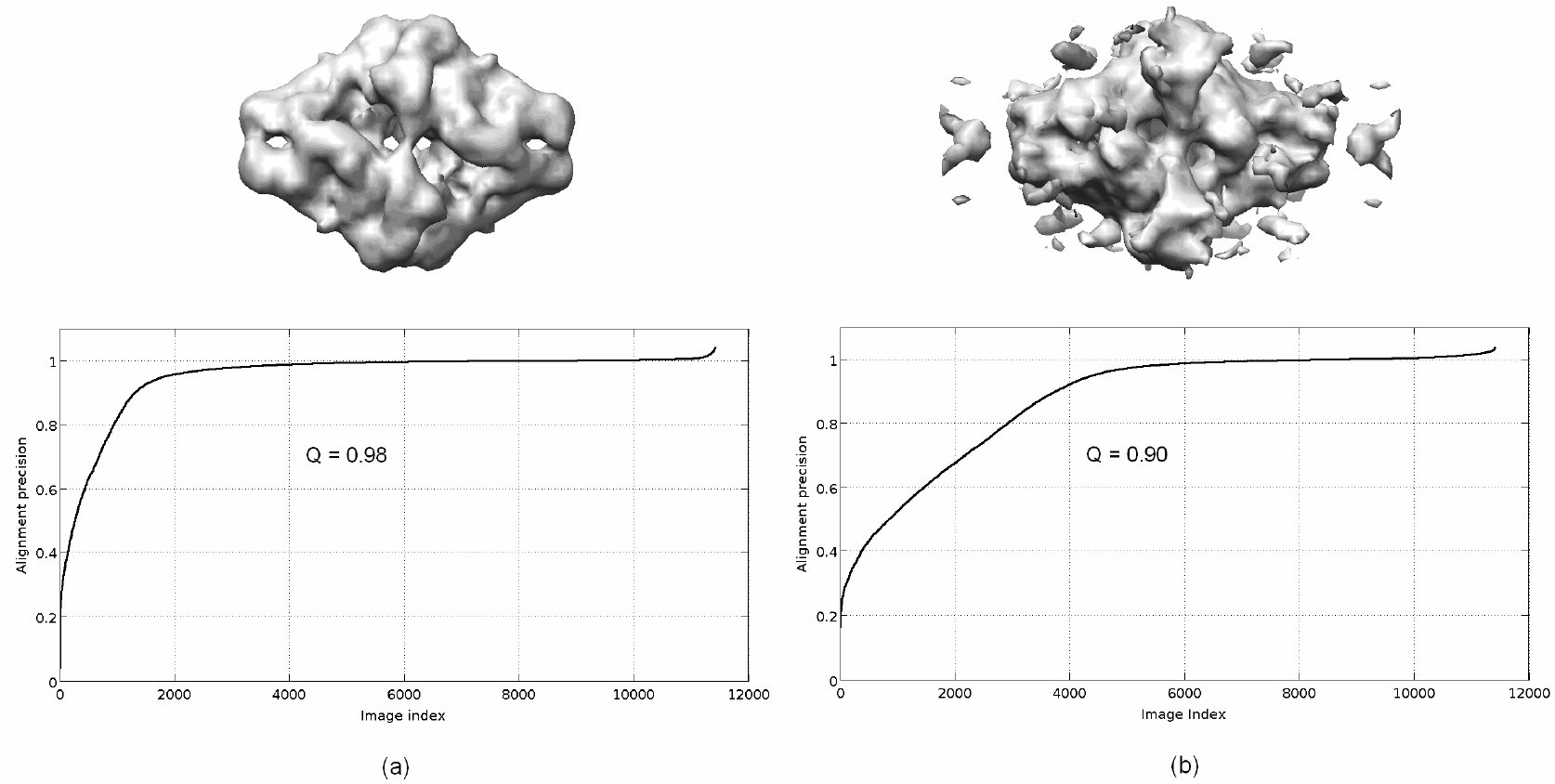
Alignment precision curves computed from the “good” (a) and “bad” quality (b) *ab initio* β-galactosidase initial volumes.

#### Third Experiment: Application to detect incorrect maps

We have applied our method to maps that have been subjected to recent controversy in the field, corresponding to reconstructions of the HIV-1 trimer with EMDB codes 5447 and 2484. We have used the experimental images previously used by the authors and deposited in EMPIAR with codes 10008 and 10004. The number of images for EMPIAR 5447 and 2484 corresponds to 124,478 and 88,125, respectively. We run our proposed soft-alignment validation approach confronting all the available particles with their respective EMDB maps. In Figure 9 we show the resulting soft-alignment validation plots for both cases. It can be visually seen that for EMPIAR 10008 (Figure 9(b)) the resultant soft-alignment plot is of low quality and most of the particles do not align with precision and accuracy. On the other hand for, EMPIAR 10004 most of the particles aligns with precision and accuracy showing a clear cluster at point (1,1). Table 2 shows the percentage of particles which aligns with precision, accuracy and both precision and accuracy in both cases. This table clearly shows that EMDB 5447 map cannot be a valid map in terms of its alignment quality.

**Figure 9.**
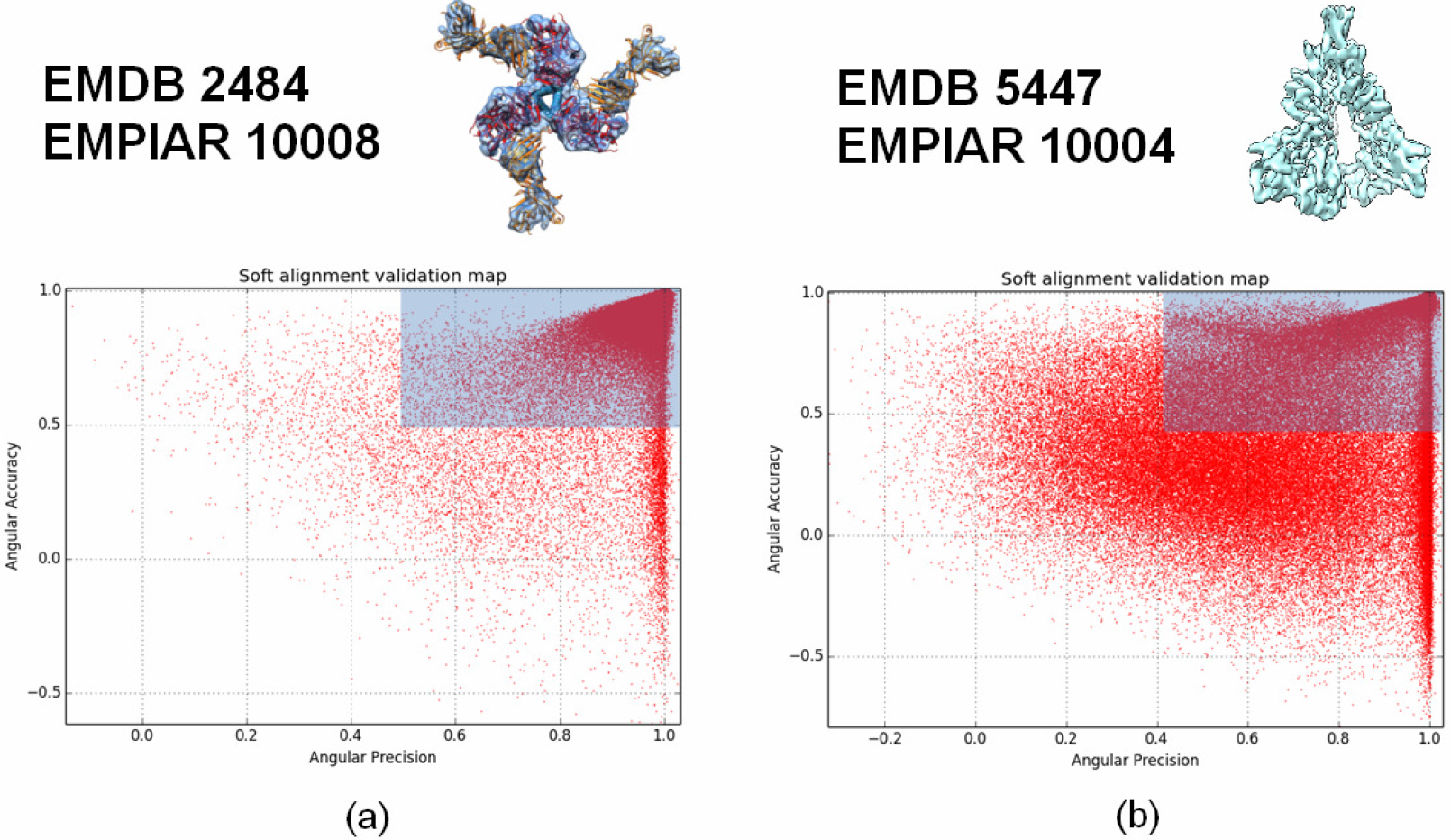
Resultant soft-alignment validation plots for HIV-1 trimmer reconstructions with EMDB codes 2484 (a) and 5447 (b) and using the deposited particles in EMPIAR with codes 10004 (88,125 particles) and 10008 (124,478 particles), respectively.

**Table 2.** Percentage of particles that align with precision, accuracy and both precision and accuracy for the HIV-1 trimmer with EMDB codes 5447 (a) and 2484 (b)

|  | **EMDB 5447** | **EMDB 2484** |
| --- | --- | --- |
| **Precision (%)** | 75 | 91 |
| **Accuracy (%)** | 45 | 98 |
| **Precision & Accuracy (%)** | 38 | 90 |

#### Fourth Experiment: Application to heterogeneous data

We have finally used our soft-alignment validation approach to data coming from the Plasmodium falciparum 80S ribosome bound to the anti-protozoan drug emetine, which is a well known heterogeneous sample. We have used the data deposited in the 2015 Map Challenge with EMPIAR ID of 10028. This data was obtained with a FEI Polara 300 microscope equipped with a Falcon II camera. The number of projection images deposited was 105,145; from this data we reconstructed a 3DEM map using Relion [Scheres2012] through Scipion framework [DelaRosa2016]. The resultant map has a 0.143-FSC resolution of 3.28 when it is compared to the PDB (PDB codes: 3j79/3j7a). The resulting soft-alignment validation plot is given in Figure 10. The percentage of particles which aligns with precision, accuracy and precision and accuracy corresponds to 0.92, 0.75 and 0.72, respectively. In this case the plot shows the highly variability of the data because the sample flexibility. Using this information we pruned particles with low alignment score so that 34% of the particles were removed. We reconstructed an additional map using the remaining 69,402 particles obtaining a 0.143-FSC resolution of 3.16 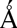 when it was compared with the PDB (PDB codes: 3j79/3j7a). In Figure 11 we show the corresponding FSC curves when the pruned and non-pruned structures were confronted with the PDB. As can be seen from Figure 11 the FSC curve obtained from the pruned data presents higher FSC values for all the frequencies, clearly indicating the better quality of this data in terms of particle homogeneity.

**Figure 10.**
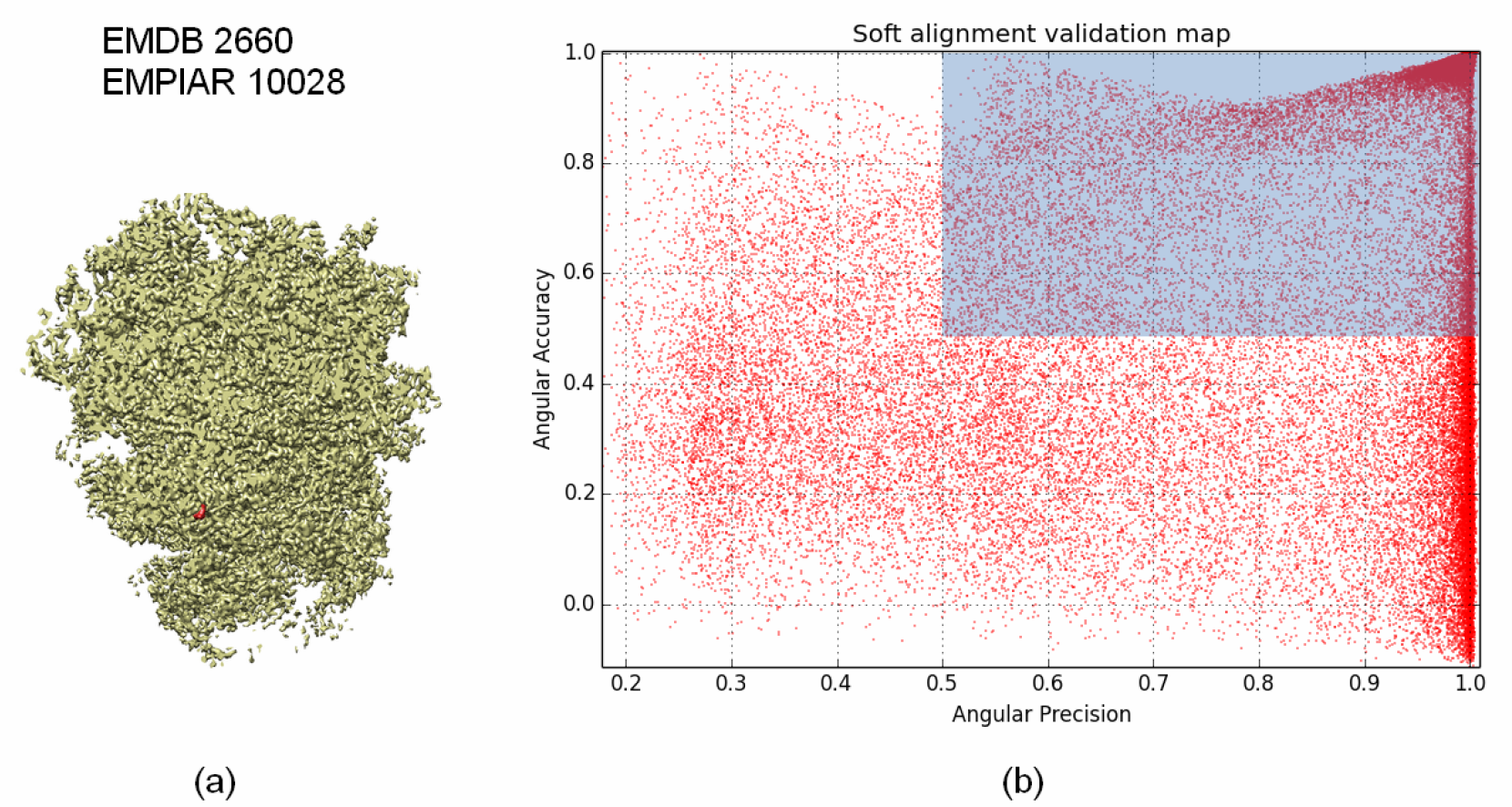
80S ribosome with EMDB code 2660 and used in the 2015 Map Challenge (a) and resultant soft-alignment validation plot (b) obtained when 69,402 particles deposited in EMPIAR 10028 were used.

**Figure 11.**
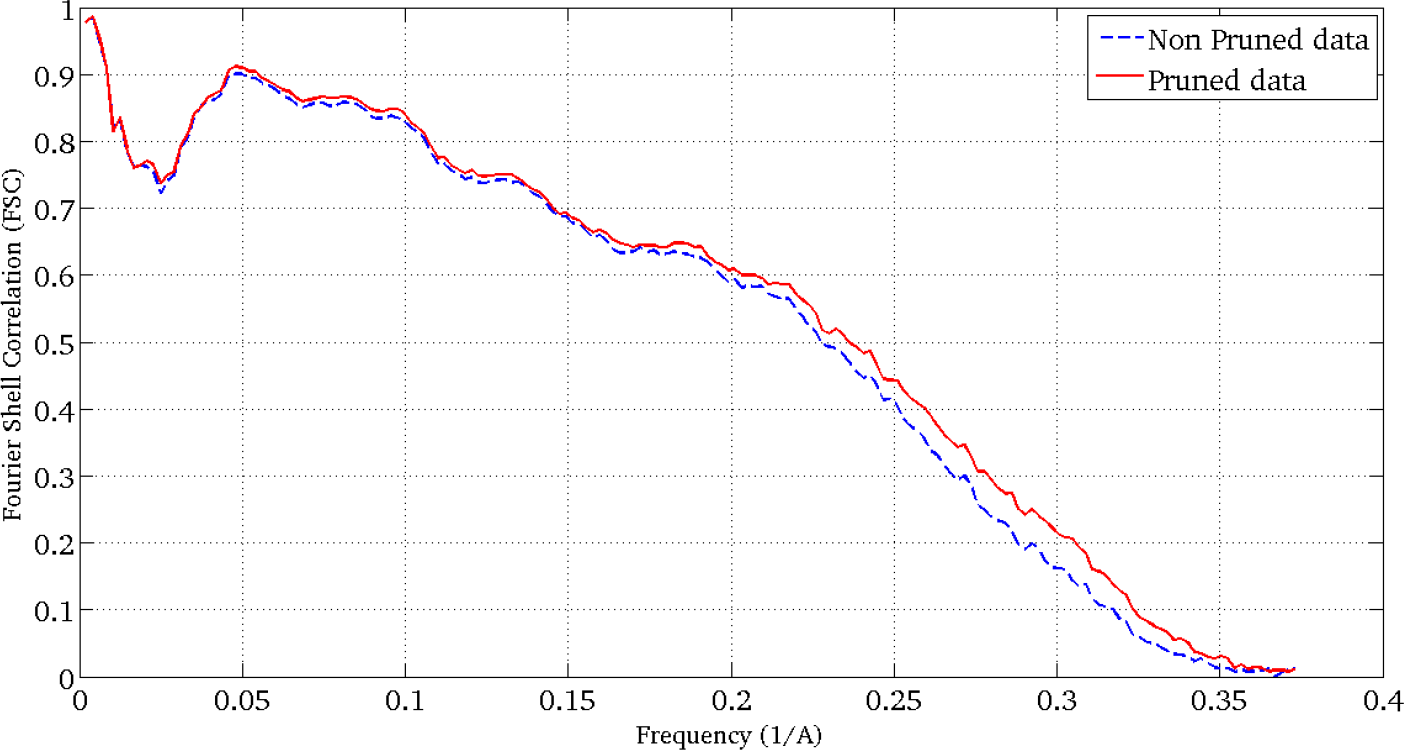
FSC curves when the pruned (red dashed curve) and non-pruned (blue solid curve) 3DEM structures were confronted with the PDB.

## Discussion

In this work we propose an approach that can determine the alignment precision and accuracy of each particle that participated in the reconstruction of a 3DEM map. Proper aligned particles should align at the same time with precision and accuracy. Usually, particles with low alignment precision score represent noise images that can not be aligned with reliability. In addition, particles with low alignment accuracy depict projections affected by artefacts or particles representing different conformations than the reconstructed 3DEM map. Pruning particles with both low alignment accuracy and precision improves the data homogeneity and therefore the quality of the reconstructed map.

The conceptual principle of the method is based on the idea that for a given map most of the experimental particles should present a cluster distribution for their most likely map orientations when a global alignment process is performed. This cluster distribution is a consequence of the spatial coherence that a correct map should show. In addition, the previously computed particle orientations, used in the map reconstruction, should be compatible with the weighted average of the most similar map orientations after the global angular search. In order to decide if a particle shows good or bad alignment precision and accuracy scores we use two references. One comes from a pure uniform random orientation distribution within the asymmetric unit. This reference exemplifies a bad alignment, where all the possible orientations have the same probability of being true. Opposed to this, we define a good reference projecting the input 3DEM map at the same orientations and in the same conditions in terms of CTF than the experimental particles but without added noise. Each experimental particle has a “perfect” counterpart which will align with precision and alignment. With these two references it is possible to define numbers between 1 and 0 describing the alignment precision and accuracy of each experimental particle. These alignment scores can be used to determine the percentage of particles that align with both accuracy and precision. We have defined as criteria that a particle aligns with precision and accuracy if these scores are equal and higher than 0.5. Using this information we can determine the percentage of particles which aligns with both precision and accuracy (*Q* value) as an indicator of the reconstruction alignment quality. Nonetheless, indicating a direct relation between the *Q* value and the map validity is difficult. It is clear that maps with low *Q* scores are affected by high heterogeneity and/or a high number of pure noise particles, among other problems, which can compromise the alignment refinement process. However, it is not possible to define a unique *Q* threshold to determine the validity of a given 3DEM map. For example, a reconstruction with a *Q* value of 0.5 can be correct if bad quality particles are just pure noise images and enough good particles are available. However, this *Q* value can be problematic if the projection images are affected by artefacts. It is clear that low values of *Q* (between 0-0.5) clearly indicate that the quality of the data is low and likely the reconstruction is not correct, and additional test should be added to show the validity of the reconstruction.

We have used our proposed method in different situations as high resolution data (β-galactosidase complex), ranking of *ab initio* initial maps (β-galactosidase complex), controversial maps (HIV-1 trimmer) and heterogeneous data (80S ribosome). In all these cases we have computed the soft-alignment validation map which gives information about the goodness of the particle alignment process and of the particle homogeneity. This information was used to correctly rank intial maps in terms of its quality and clearly discard the HIV-1 trimmer map in terms of its alignment quality. In addition, these soft-alignment validation plots were used to improve the quality of the data in terms of its homogeneity by a pruning process. After rejecting particles with low alignment scored we clearly improved the resolution of the reconstructed maps. In the case of the β-galactosidase and 80S ribosome macromolecules we improved from 3.02 and 3.28 to 3.00 and 3.16 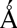, respectively.

